## Supplementary Material for "Gain-of-function dynamin-2 mutations linked to centronuclear myopathy impair Ca^2+^-induced exocytosis in human myoblasts"

Index

**Figure S1**: $Ca^{2+}$ signals induced with ionomycin at 1, 5 and 10 µM.

**Figure S2**: Non-lateral and lateral diffusion event detection.

**Figure S3**: Effects of an acidic extracellular solution on neutral vesicles.

**Figure S4**: Effects of 100 mM HEPES on decay time.

**Figure S5**: Percentage of transfected cells WT dynamin-2 or the mutants A618T or S619L.

**Figure S6**: $Ca^{2+}$ signals induced with 10 µM ionomycin in C25 myoblasts expressing WT dynamin-2, or the p.A618T or p.S619L mutations.

**Figure S7**: Comparison of the $Ca^{2+}$ and pHluorin signals of C25 myoblasts non-transfected and transfected with WT dynamin-2.

**Figure S8**: Non-lateral and lateral diffusion fluorescence events in a C25 myoblasts expressing the dynamin-2 mutation p.A618T.

**Figure S9**: Ionomycin stimulation promotes GLUT4 translocation from intracellular stores to the surface membranes of human myoblasts.

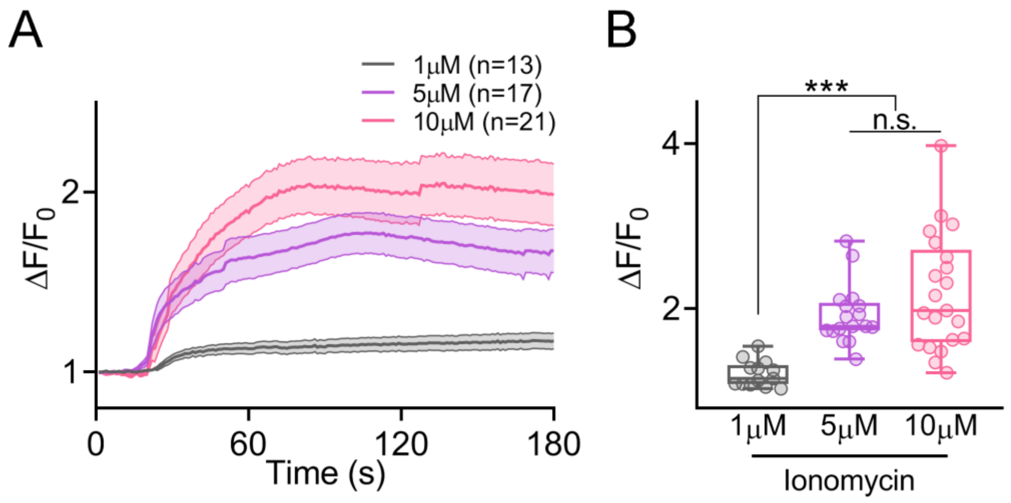


**Figure S1**: *$Ca^{2+}$ signals induced with ionomycin at 1, 5 and 10 µM*. Cytosolic $Ca^{2+}$ signals induced with different concentrations of ionomycin were recorded in C25 myoblasts bathed in a solution containing 2 mM $Ca^{2+}$, and analyzed by epifluorescence time lapse imaging. **A**: Time-lapse images of the $[Ca^{2+}]_i$ transients evoked by application of ionomycin are shown as relative changes in fluorescence intensity ($\Delta F/F_0$). Data represent mean ± SEM. **B**: Each dot represents $\Delta F/F_0$ maximum values from individual cells from three different cultures. Horizontal lines indicate medians and whiskers max and min values of the distribution. The number of cells analyzed is indicated in parentheses in Panel A. Experimental details are given in Materials and Methods. ***$p<0.001$ (Kruskal-Wallis test followed by Dunn's multiple comparison test).

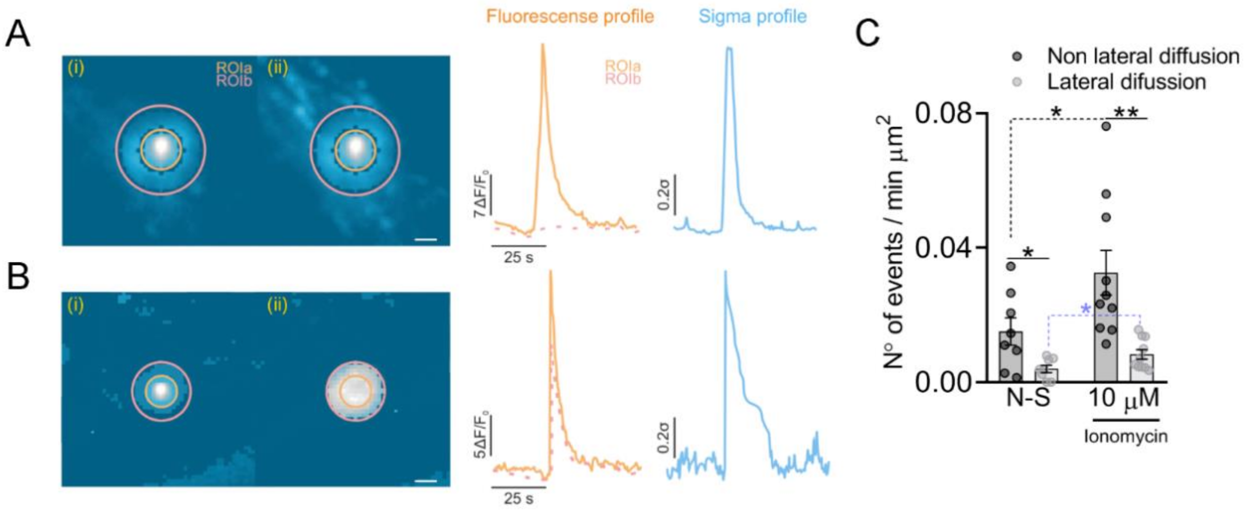


**Figure S2**: *Non-lateral and lateral diffusion event detection.* Representative non-lateral (A) and lateral (B) diffusion events are displayed with their corresponding normalized fluorescence and sigma profile in time (see Methods). Thermal panels (i) show the fluorescence of one representative event in its first frame of appearance (t=0), while thermal panels (ii) show all maximum values of fluorescence, from all consecutive frames of the same event, overlapped. Orange and pink circles indicate the regions of interest (ROI) ROIa and ROIb, respectively. Scale bar = 0.5 µm. (C) Quantification of the number of non-lateral (black) and lateral (light grey) diffusion events, normalized by area and time, in non-stimulated cells (N-S, n=8) and stimulated with 10 µM of ionomycin (10 µM, n=10). Each dot represents individual cells obtained from at least three different cultures. Bars indicates means with SEM. Black solid lines indicates statistical comparison between the number of non-lateral and lateral diffusion events within each experimental group (N-S, 10 µM), while dashed black and blue lines indicates statistical comparison of the number of non-lateral and lateral diffusion events between different experimental groups, respectively. *p < 0.05 and **p < 0.025 (Student's t-test).

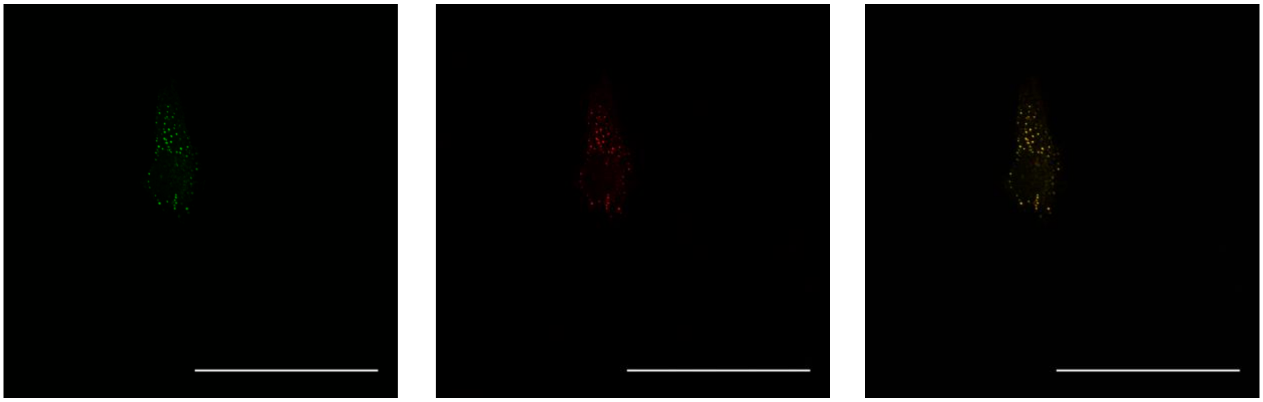


**Figure S3**: *Effects of an acidic extracellular solution on neutral vesicles.* To know whether an acidic extracellular solution (pH=5.5) quenches the intravesicular IRAP-pHluorin, C25 cells were visualized in a confocal microscopy using a C1 Plus laser-scanning confocal microscope (Nikon, Japan) and a Nikon Fluor 60x/1.00 W objective (Nikon, Japan). The panel shows example of confocal images acquired in cells kept in pH 7.4 extracellular solution (left panel; pseudocolor: green) and after 50 s in an acidic extracellular solution (middle panel; pseudocolor: red) and the merge of both images (pseudocolor: orange). Scale bar = 100 μm. The brightness was increased to 14% to better visualize the fluorescence of the vesicles containing IRAP-pHluorin. The analyses of 5 cells yielded Pearson's correlation coefficients of 0.85 ± 0.03. The composition of the acidic solution was (in mM): 140 NaCl, 2.4 KCl, 2 $CaCl_2$, 2 $MgCl_2$, 10 glucose, 10 HEPES and 10 citric acid (pH=5.5).

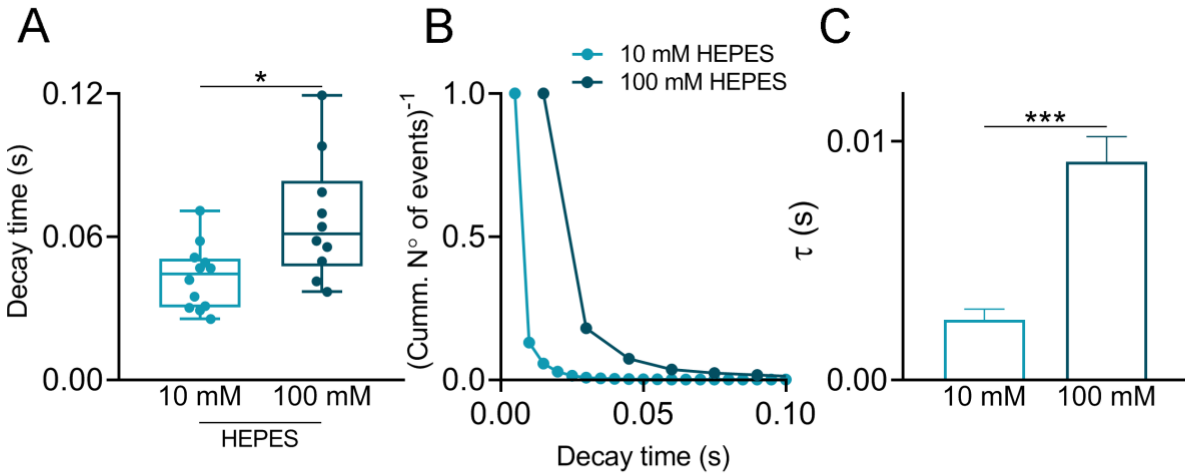


**Figure S4**: *Effects of 100 mM HEPES on decay time*. **A**: Comparison of averaged decay times of the fluorescence events induced with 1 µM ionomycin in an extracellular solution containing 10 o 100 mM HEPES (12 and 10 cells, respectively). For the experiments with 100 mM HEPES, the NaCl concentration was adjusted to keep osmolarity constant. Dots represent values from individual cells from at least three independent cultures. *$p<0.05$ (Student's test). **B**: Survival curves (inverse of the normalized cumulative distributions) in both conditions fit with single exponential decay function ($R^2$ = 0.991 and 0.996 for 10 and 100 mM HEPES, respectively). **C**: Time constants ($\tau$) (media ± SE) obtained from their fitting to a single exponential decay function. Comparisons of both conditions were performed by Student's t-test with Welch's correction, ***$p<0.001$.

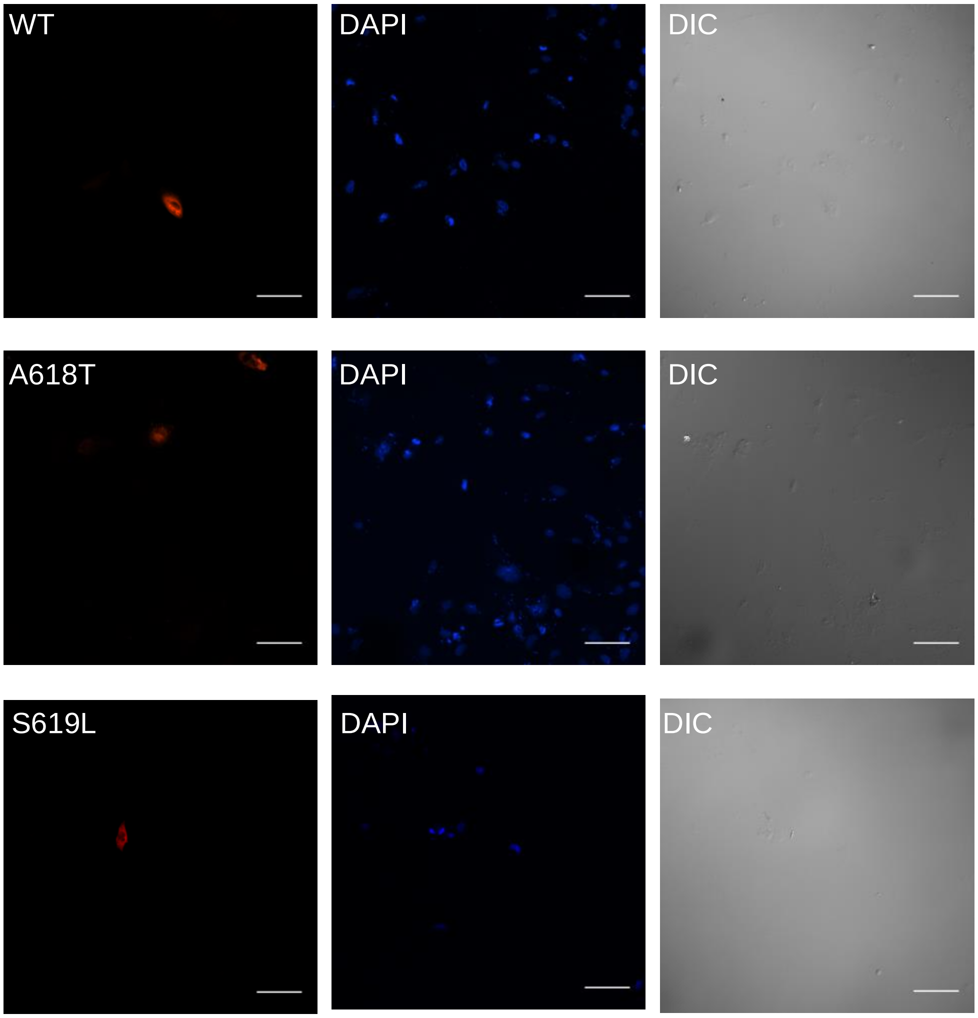


**Figure S5**: *Percentage of transfected cells with WT dynamin-2 or the mutants A618T or S619L*. To determine the percentage of cells transfected with the dynamin-2 variants, cells were fixed with 4% p-formaldehyde 24 h after transfection, stained with 5 mg/mL 4,6-diamidino-2-phenylindole (DAPI) and visualized at the equatorial plane in a C1 Plus laser-scanning confocal microscope (Nikon, Japan) using a 40× objective. The panels show examples of confocal images of C25 cells expressing WT dynamin-2 or the mutants A618T or S619L. The analyses of 13 coverslips from three different cultures yielded percentage of transfected cells of 2.2 ± 0.7, 3.5 ± 0.7 and 2.3 ± 0.7 for WT dynamin-2 or the mutants A618T or S619L, respectively.

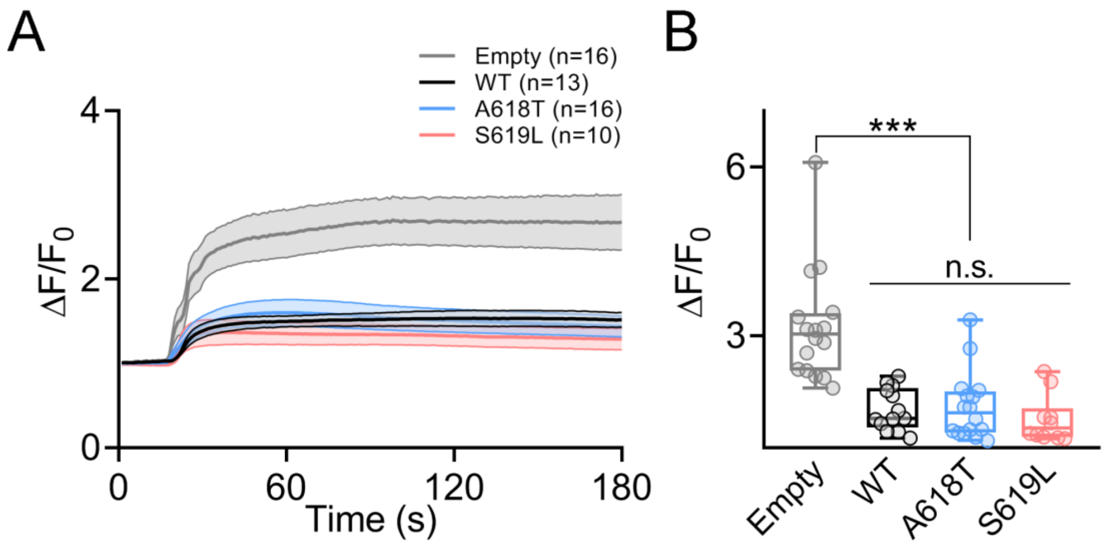


**Figure S6**: *Cytosolic $Ca^{2+}$ signals induced with 10 μM ionomycin in C25 myoblasts expressing the empty mCherry vector, WT dynamin-2 or p.A618T and p.S619L mutants* were recorded and analyzed by epifluorescence time lapse imaging. **A**: Ionomycin-evoked $[Ca^{2+}]_c$ transients expressed as relative changes in fluorescence intensity ($\Delta F/F_0$) obtained from time-lapse images. Data represent mean ± SEM. **B**: Each dot represents $\Delta F/F_0$ maximum values from individual cells from three different cultures. Horizontal lines indicate medians and whiskers max and min values of the distribution. The number of cells analyzed is indicated in parentheses in Panel A. ***$p<0.001$ compared with the empty vector. Non-significant differences were found between WT dynamin-2 and the p.A618T and p.S619L mutants (Kruskal-Wallis test followed by Dunn's multiple comparison test). Experimental details are given in Materials and Methods.

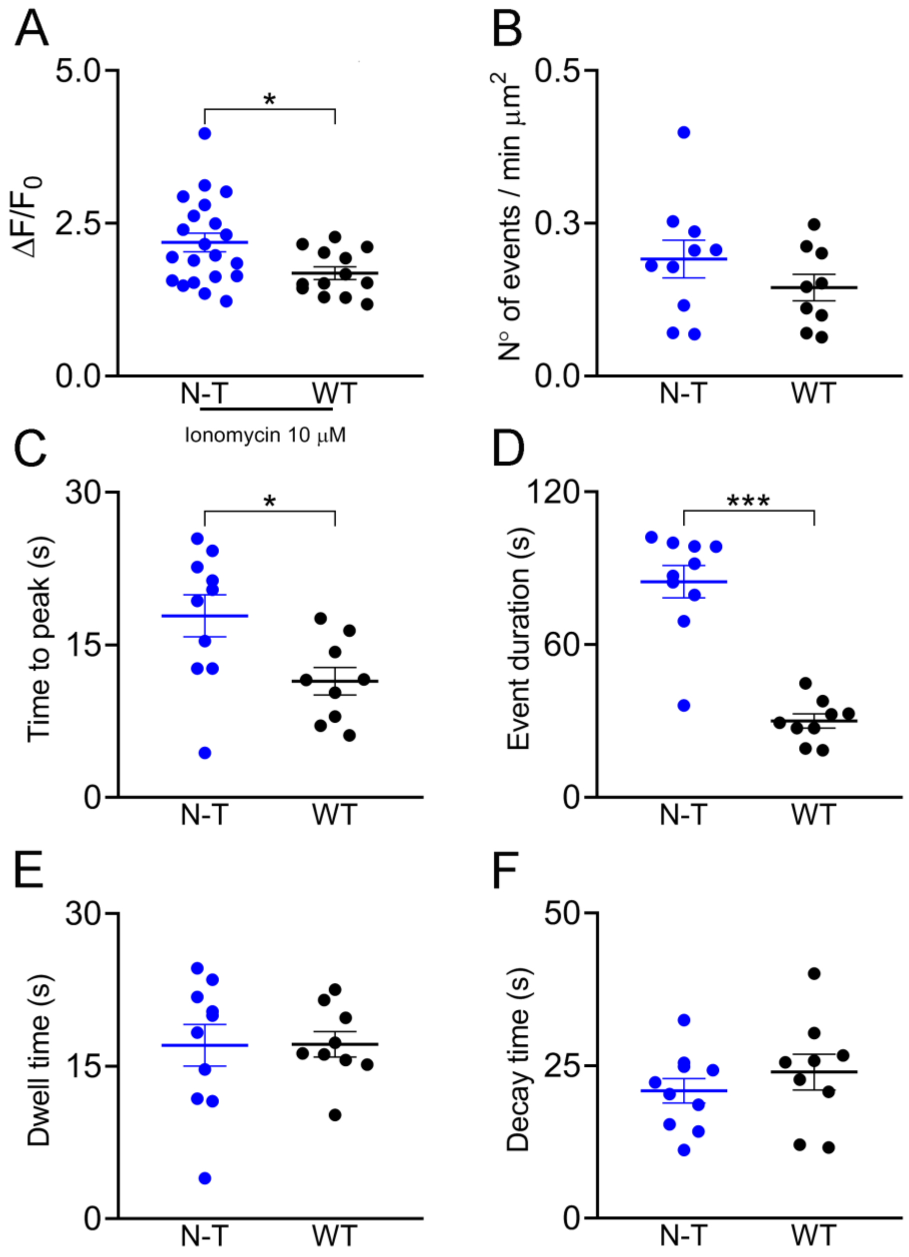


**Figure S7**: *Comparison of the $Ca^{2+}$ and pHluorin signals of C25 myoblasts non-transfected (N-T) and transfected with WT dynamin-2 (WT).* **A**: $Ca^{2+}$ signal amplitudes. **B**: Event frequencies. **C**: Times to peak. **D**: Event durations. **E**: Dwell-times. **F**: Decay times. Each dot represents an individual cell and at least 10 cells for each condition were recorded from at least three different cultures. Horizontal lines indicate means with SEM. *p<0.05, ***p < 0.001 (Student's t-test).

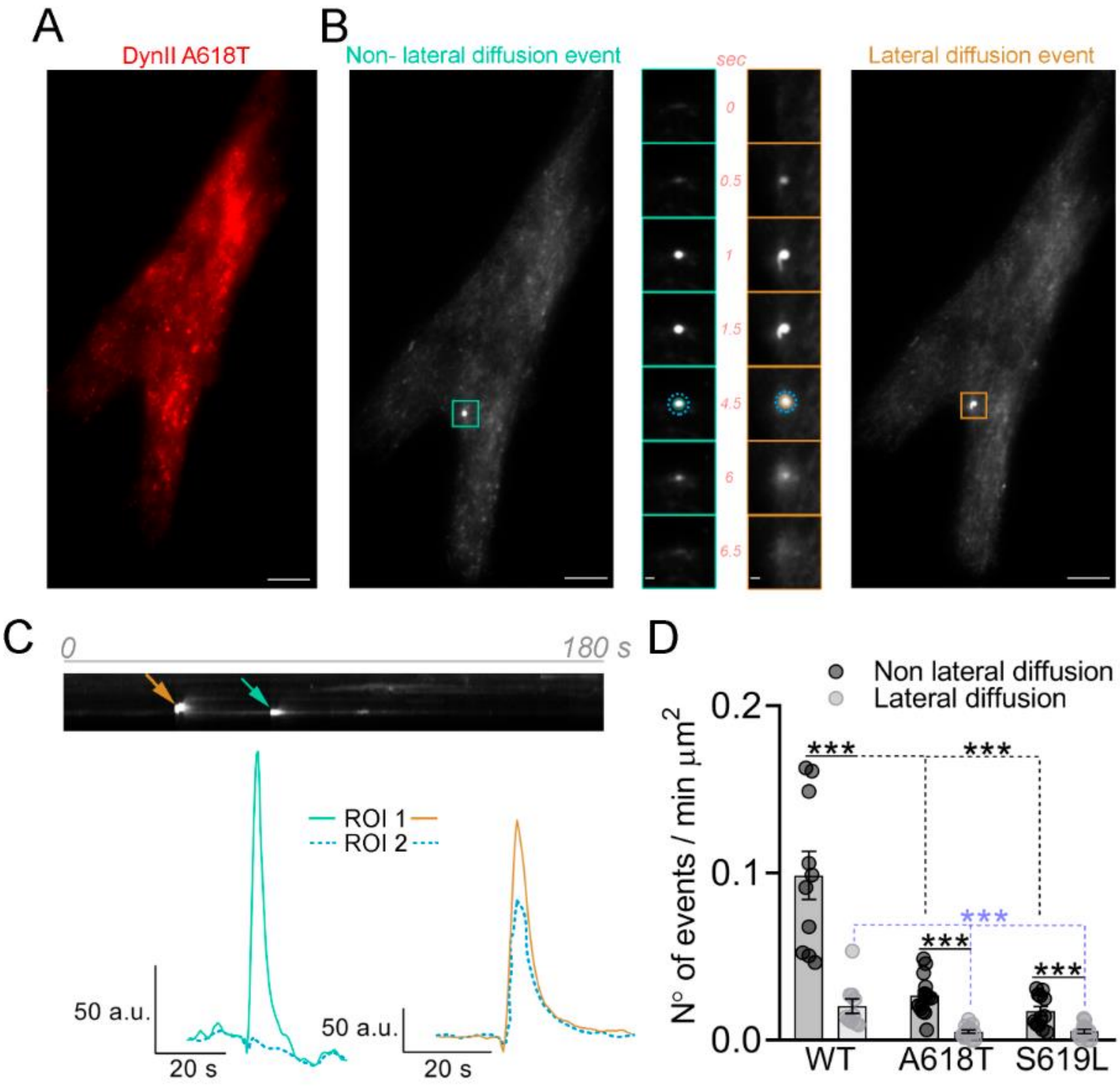


**Figure S8**. *Non-lateral and lateral diffusion events quantification*. **A**: Epifluorescence image of a C25 cell expressing the dynamin-2 mutation p.A618T fused to the mCherry reporter. Scale bar = 20 µm **B:** The same C25 cell but visualized in the TIRFM plane displaying a non-lateral (left, turquoise square) and a lateral diffusion (right, orange square) events. The middle panels in B show the sequences of video frames of those events (the time 0 corresponds to the show-up of the fluorescence spot); the solid and dashed line circles at 4.5 seconds represent ROI 1 (25 x 25 pixels) and ROI 2 (50 x 50 pixels), respectively. Scale bar in middle panels is 0.5 µm. Note that the non-lateral diffusion event vanished without diffusing outside the ROI 1, whereas the lateral diffusion event spread outside the ROI 1. **C**: The upper panel shows the kymograph corresponding to the selected areas (the time corresponds to the whole duration of the recording period). The turquoise and orange arrows indicate the non-lateral and lateral diffusion events, respectively. The lower panel shows fluorescence intensity profiles inside the ROI 1 (solid lines) and in the surrounding area (ROI 2, dash lines) (see Methods). **D:** Quantification of the number of non-lateral (dark gray) and lateral (light grey) diffusion events, normalized by area and time, in cells expressing dynamin-2 WT (n=10) and the dinamyn-2 mutations p.A618T (n=14) and p.S619L (n=12). Quantification of both types of events were performed by the analysis of the evolution of the sigma parameter in the time, as described in Methods. Each dot represents individual cells obtained from at least three different cultures. Bars indicates means with SEM. Black solid lines indicates statistical comparison between the number of non-lateral and lateral diffusion events within each experimental group (WT, A618T, S619L), while dashed black and blue lines indicates statistical comparison of the number of non-lateral and lateral diffusion events between different experimental groups, respectively. ***$p < 0.001$ (Student's t-test).

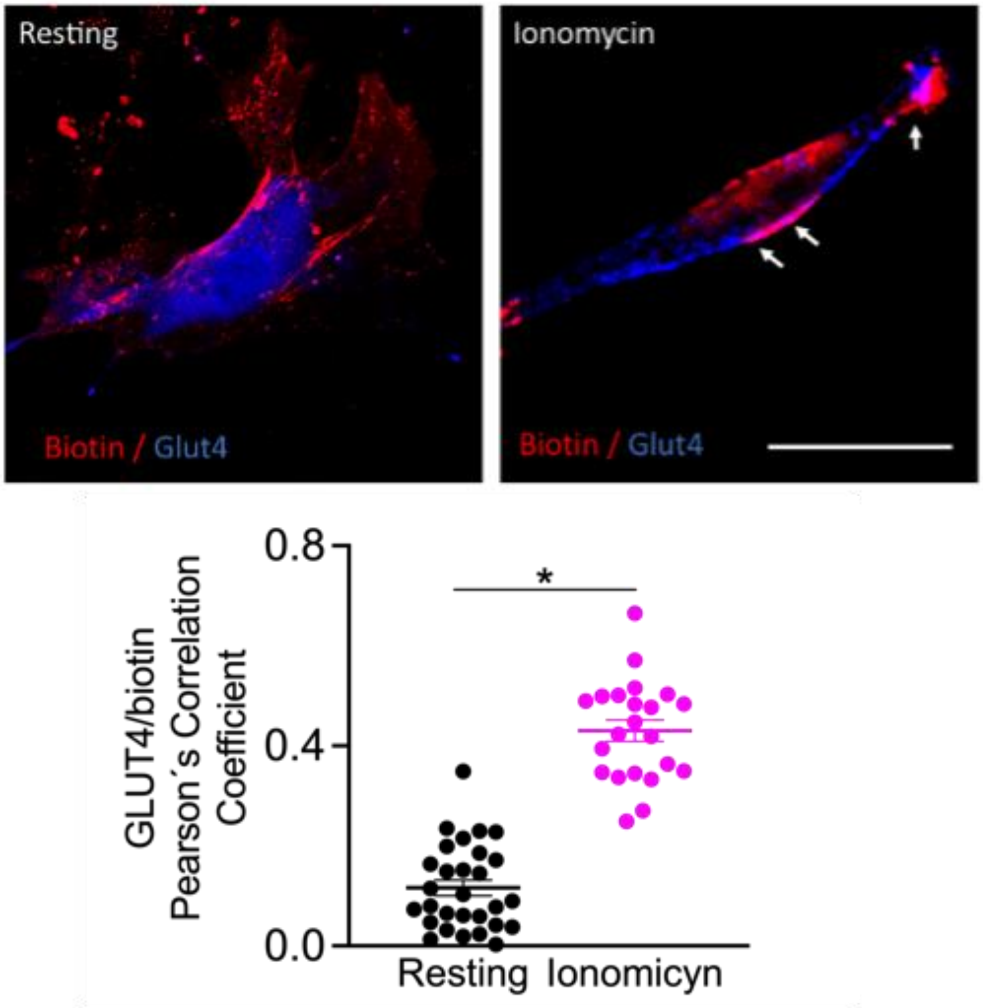


**Figure S9**. *Ionomycin stimulation promotes GLUT4 translocation from intracellular stores to the surface membranes of myoblasts.* RCMH human myoblasts were subjected to biotinylation of the surface proteins at 4ºC for 30 min and stimulated with 20 µM ionomycin during 30 s to induce endogenous GLUT4 translocation from intracellular stores to the surface membrane. RCMH myoblast were fixed with 4% PFA, immunolabeled with an antibody against GLUT4 (blue) and biotinylated proteins were stained with streptavidin-Cy3 (red) and visualized by confocal microscopy. Note that ionomycin stimulation significantly increases colocalization between GLUT4 and biotinylated proteins (estimated by the Pearson`s correlation coefficient), indicative of GLUT4 insertion into the surface membrane. White arrows indicate colocalization sites in the membrane of myoblasts. Bars are mean ± SEM. *$p<0.05$ compared to resting condition. (29 resting cells and 22 ionomycin stimulated cells).
